# Alterations in Circulating T-Cell Subsets with Gut-Homing/Residency Phenotypes Predict HIV-1 Status and Subclinical Atherosclerosis

**DOI:** 10.1101/2024.11.22.624885

**Authors:** Etiene Moreira Gabriel, Jonathan Dias, Abdelali Filali-Mouhim, Ramon Edwin Caballero, Tomas Raul Wiche Salinas, Manon Nayrac, Carl Chartrand-Lefebvre, Jean-Pierre Routy, Madeleine Durand, Mohamed El-Far, Cécile Tremblay, Petronela Ancuta

## Abstract

Antiretroviral therapy (ART) controls HIV-1 replication in people with HIV-1 (PWH), but immunological restauration is not achieved at mucosal barriers. Intestinal integrity impairment fuels microbial translocation and chronic immune activation, thus heightening the cardiovascular disease (CVD) risk. Here, we sought to identify novel immunological predictors of the HIV and CVD status in the peripheral blood of ART-treated PWH (HIV^+^; n=42) and uninfected participants (HIV-; n=40) of the Canadian HIV and Aging Cohort Study (CHACS), with/without subclinical coronary atherosclerotic plaques, measured by Coronary Computed Tomography Angiography as total plaque volume (TPV, mm^3^). PBMCs were analyzed by flow cytometry for the expression of T-cell lineage (CD45, CD3, CD4, CD8αα, CD8αβ, TCRαβ, TCRγδ), epithelial cell (EpCAM/CD326), activation (HLA-DR), and gut-homing/residency markers (CD69, CD196/CCR6, CD199/CCR9, CD49d/Itgα4, CD103/ItgαE, Itgβ7). CellEngine supervised clustering of the flow cytometry data revealed profound alterations in the CD3^+^ T-cell pool in relationship with the HIV status, with the accumulation of peculiar CD8^+^TCRαβ^+^ and TCRγδ^+^ cells, to the detriment of CD4^+^TCRαβ^+^ subsets. FlowJo manual gating further identified CD4^+^ T-cell subsets with peculiar CD326^+^CD69^+^CCR6^+^ItgαE^+^ and CCR6^+^Itgβ7^-^ phenotypes that were increased in frequency in HIV^+^ *versus* HIV^-^ participants, together with a decreased frequency of CD8^+^ T-cells with an intraepithelial lymphocyte (IEL)-like CD3^+^CD4^-^ TCRαβ^+^TCRγδ^-^CD8αα^+^CD8αβ^-^ phenotype. Finally, multivariate logistic regression revealed the predictive capacity of specific T-cell subsets regarding the HIV/CVD status, and TPV values. Of particular relevance, we identified ItgαE^+^CD8^+^, ItgαE^-^CD8^+^, CCR6^+^CD4^+^, and CCR6^+^Itgβ7^-^ CD4^+^ T-cell subsets as strong positive predictors of atherosclerotic plaque volume in crude models or upon adjustment for HIV/CVD confounding factors.

## INTRODUCTION

HIV-1 affects an estimated 39.0 million individuals globally, with approximately 1.3 million new incidences reported in 2023 [1]. While antiretroviral therapy (ART) has significantly improved the lives of people with HIV-1 (PWH), transforming it into a manageable chronic condition, treatment is not curative [2–5]. Consequently, ART-treated PWH experience premature aging and increased risk of developing non-AIDS comorbidities, such as cardiovascular disease (CVD) [6–8]. Therefore, new efforts are required to find solutions for the current medical challenges faced by PWH, even in countries where ART is available.

HIV-1 infection provokes profound immunological alterations leading to inadequate mucosal barrier functions. This can result in cells displaying modified immune phenotypes related to inflammation and cell activation [9, 10]. For example, the expression of HLA-DR is identified as an indicator of overt immune activation associated with HIV-1 disease progression [11, 12], while the expression of CD69, an early T-cell activation marker [13], indicates the ability of cells to reside into tissues [14]. The gut-associated lymphoid tissues (GALT) represent the primary location for CD4^+^ T-cell depletion during the acute phase of HIV-1 infection and also constitute an important site for HIV-1 reservoir persistence during the chronic phase [3–5, 15–17]. ART initiation promptly decreases HIV-1 viral loads and results in the normalization of CD4^+^ T-cell counts in the bloodstream. However, even in cases where viral suppression occurs and circulating CD4^+^ T-cell counts are restored, PWH continue to exhibit elevated CD8^+^ T-cell counts, resulting in persistently imbalanced CD4/CD8 ratios, with a low CD4/CD8 ratio predicting the size of HIV reservoirs [18] and morbidity/mortality [19, 20]. In addition, the replenishment of the CD4^+^ T-cell pools within the GALT progresses slowly and is less complete than in the peripheral blood [21]. This inadequate recovery of GALT-infiltrating CD4^+^ T-cells leads to a breach in the epithelial cell barrier, promoting microbial translocation and triggering immune activation [15, 22]. These events, uncontrolled by ART, are documented to fuel a state of chronic inflammation and immune activation facilitating the occurrence of non-AIDS comorbidities in ART-treated PWH, including premature immunological aging and CVD [6–9, 23–25]. Effective communication between intestinal epithelial cells (IEC) and immune cells is crucial for maintaining mucosal homeostasis in its capacity to fight pathogens at barrier surfaces [26, 27]. Th17-polarized CD4^+^ T-cells are fundamental for sustaining gut immunity, significantly aiding in maintaining epithelial integrity and functionality [28–31]. Consequently, the deficit in Th17 cells profoundly alters the GALT barrier functions, leading to a compromised epithelium and chronic systemic inflammation [9, 28–30, 32]. Pioneering studies documented that HIV-1 infection is associated with changes in the expression of the gut-homing molecule integrin (Itg)α4β7 on the peripheral CD4^+^ T-cells [33–35] and that the expression of the Itgα4β7 on blood CD4^+^ T-cells mirrors events taking place in the gut and predicts the risk of HIV acquisition/disease progression [36]. Since the Itgα4β7^+^ CD4^+^ T-cell subset predominantly consists of CCR6^+^ Th17 cells, their capacity to migrate into the GALT justifies our findings that CCR6^+^Itgβ7^+^ CD4^+^ T-cells are selectively targeted by HIV-1 for infection [28–30, 33]. Indeed, Itgα4β7 has been reported as a marker of CD4^+^ T-cells highly permissive to HIV-1 infection [33]. In addition, we demonstrated that retinoic acid promoted Itgβ7 expression in CCR6^+^ CD4^+^ T-cells, and increased HIV-1 replication and outgrowth [28–30]. More evidences underscore how cellular immune activation is linked to the migration and subsequent depletion of Th17 cells in the gut of PWH *via* CCR6/CCL20 and CCR9/CCL25-dependent mechanisms[12, 15, 37–39]. In addition to Th17 cells, other T-cell subsets important for the maintenance of gut immunity include the intraepithelial lymphocytes (IELs), a subset marked by the expression of CD103/ItgαE [40, 41]. Importantly, IELs are tissue-resident T-cells that usually do not recirculate in the blood [40, 42–46], with functional alterations in IELs being reported during HIV-1 infection [47, 48].

Considering this knowledge, we hypothesized that alterations in the frequency and phenotype of circulating T-cells, as a consequence of an impaired intestinal barrier integrity, contribute to an increased risk of coronary atherosclerosis occurrence in ART-treated PWH. In an effort to identify a novel blood immunological signature indicative of HIV status and CVD risk, in this study, we had access to PBMC samples from ART-treated PWH and HIV-uninfected controls included in the Canadian HIV and Aging Cohort Study (CHACS), with and without subclinical atherosclerosis, determined by Computed Coronary Tomography Angiography (CCTA) measurement of the coronary artery atherosclerotic, as we previously reported [49, 50]. Polychromatic flow cytometry analysis was performed using a large panel of T-cell lineage (CD45, CD3, CD4, CD8αα, CD8αβ, TCRαβ, TCRγδ), epithelial cell (EpCAM/CD326), activation (HLA-DR), and gut-homing/residency markers (CD69, CD196/CCR6, CD199/CCR9, CD49d/Itgα4, CD103/ItgαE, Itgβ7). This allowed an in depth characterization of circulating T-cell subsets differentially expressed in HIV^+^ *versus* HIV^-^ CHACS participants. Multivariate logistic regression analysis explored the relationship between the identified T-cell subsets and the HIV and CVD status, as well as total plaque volume (TPV) values, in crude models and upon adjustment for confounding factors. Together, our findings demonstrate that alterations in circulating T-cell subsets with gut-homing/residency phenotypes predict the HIV status and the presence of subclinical atherosclerotic plaque in ART-treated PWH.

## MATERIAL AND METHODS

### Study Participants

The Canadian HIV and Aging Cohort Study (CHACS) has been described in previous publications by our group [49–53]. In summary, the inclusion criteria for the cardiovascular imaging subset of the CHACS cohort required participants to be over 40 years old, have a 10-year CVD risk as determined by the Framingham Risk Score (FRS – which calculates risk based on factors such as age, cholesterol levels, systolic blood pressure, smoking status, and diabetes status) ranging from 5-20%, and have no clinical diagnosis of CVD. Participants with renal impairment or a known hypersensitivity to contrast agents were excluded from the study. Here, we used PBMCs from 82 CHACS participants, of whom n=42 were ART-treated PWH (HIV+ART or HIV^+^) and n=40 were HIV-uninfected participants (HIV^-^). The clinical parameters of the study participants are depicted in Tables 1-2. Further stratification of both groups was conducted based on the presence or absence of subclinical atherosclerotic plaque on CCTA, determined by TPV, measured in cubic millimeters (mm^3^ equal to or greater than zero), as we previously reported [49–53].

**Table 1:**
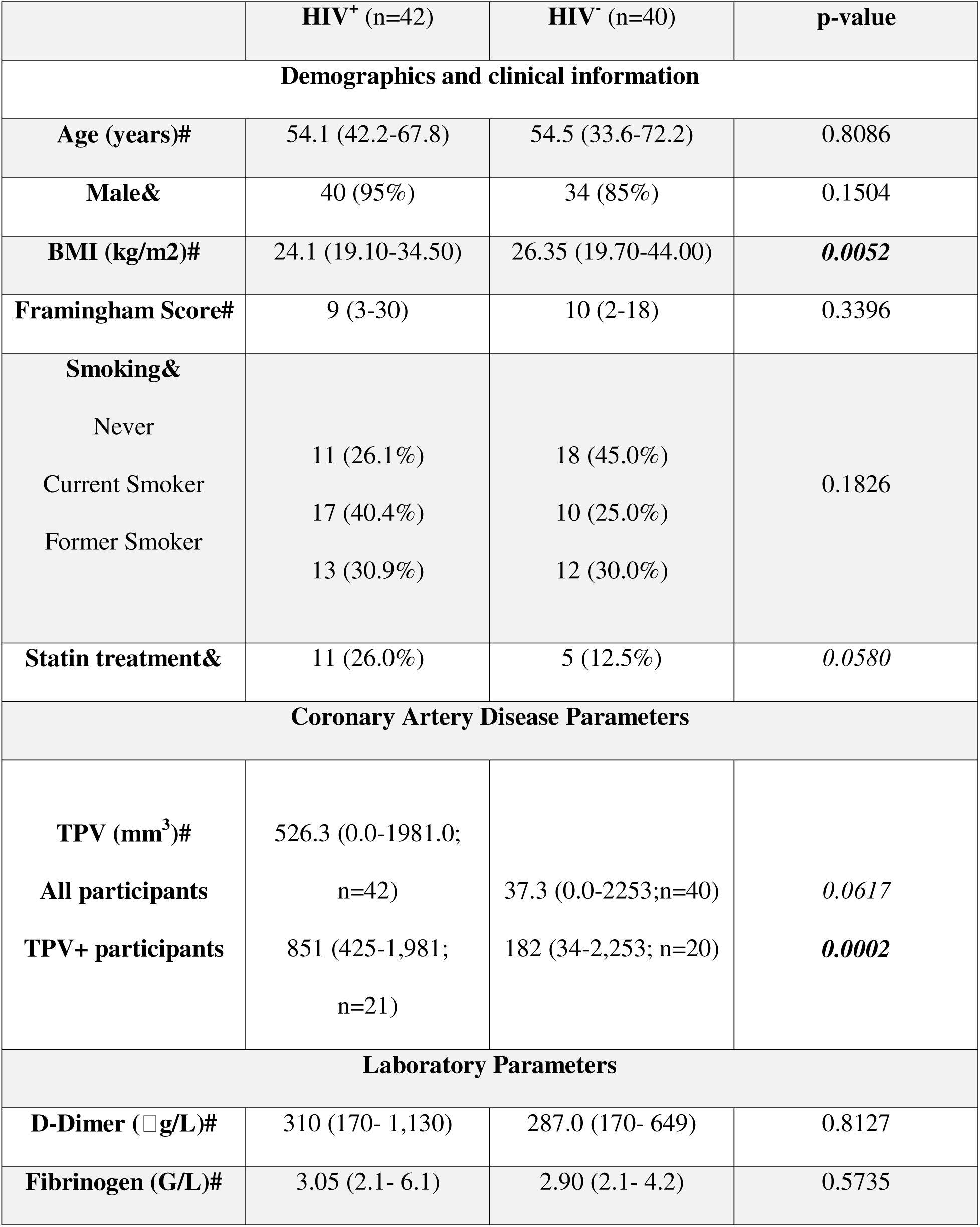
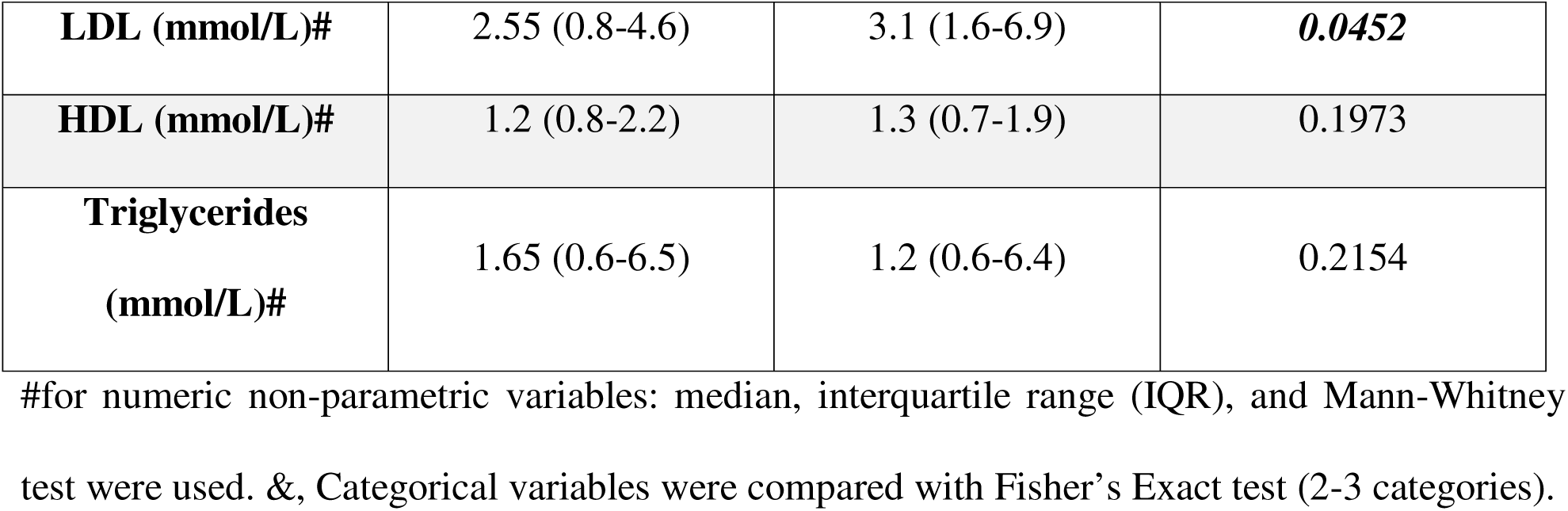
Description of study CHACS participants

**Table 2:** Clinical parameters of ART-treated PWH CHACS participants with and without subclinical atherosclerosis

|  | TPV+ (n=24) | TPV- (n= 16) | p-value |
| --- | --- | --- | --- |
| **Demographics and clinical information** |  |  |  |
| **Age (years)#** | 53.9 (42.2-64.1) | 54.3 (42.2-62.1) | 0.6573 |
| **Male&** | 23 (95.8%) | 15 (93.7%) | >0.9999 |
| **BMI (kg/m2)#** | 22.9 (19.1-34.5) | 25.5 (21.0-28.2) | 0.1320 |
| **Framingham Score#** | 9.0 (4-30) | 8 (3-18) | 0.1404 |
| **Smoking&** |  |  |  |
| Never Smoked | 4 (16%) | 5 (31%) | 0.0666 |
| Current Smoker | 14 (58%) | 3 (18%) |  |
| Former Smoker | 6 (25%) | 7 (43%) |  |
| N/A | 0 (0%) | 1 (6%) |  |
| **Statin treatment&** | 6 (25%) | 5 (31%) | 0.7275 |
| **Laboratory Parameters** |  |  |  |
| **D-Dimer (ug/L)#** | 340 (170- 720) | 275.0 (170- 1130) | 0.3029 |
| **Fibrinogen (G/L)#** | 3.2 (2.1- 6.1) | 2.8 (2.1- 3.7) | 0.0827 |
| **LDL (mmol/L)#** | 2.4 (0.8- 4.3) | 2.8 (1.9- 4.6) | 0.2972 |
| **HDL (mmol/L)#** | 1.1 (0.8-2.2) | 1.2 (0.8-2.2) | 0.7596 |
| **Triglycerides**<br>**(mmol/L)#** | 1.7 (0.6-6.5) | 1.5 (0.8- 3.2) | 0.2033 |
| **Serology** |  |  |  |
| **Co-infection CMV#** | 20 (83%) | 12 (75%) | 0.5227 |

| HIV disease parameters |  |  |  |
| --- | --- | --- | --- |
| **Duration of HIV**<br>**(years)#** | 20.6 (5.7-30.8) | 17.7 (3.8-29.4) | 0.2363 |
| **Duration of ART**<br>**(years)#** | 16.6 (3.3-24.8) | 13.6 (0.0-25.0) | 0.0706 |
| **Nadir CD4 (x10<sup>9</sup>/L)#** | 235 (20-556) | 190 (10-750) | 0.7889 |
| **CD4 (%)#** | 31.0 (3-46) | 35.0(13-57) | 0.2250 |

### Ethical statement

The collection of peripheral blood from the participants was conducted in compliance with the Declaration of Helsinki. This research received approval from the Centre de Recherche du CHUM, Montréal, Québec, Canada (#CE.11.063). Written informed consents were obtained from all participants.

### Blood collection and phenotypic analysis

Peripheral blood mononucleated cells (PBMCs) were isolated by Ficoll density centrifugation from peripheral blood collected by venipuncture on EDTA Vacutainer tubes. PBMCs were frozen in DMSO 10% FBS until use. Thawed PBMCs were stained with the following fluorochrome-conjugated antibodies: CD3 (AF700), CD45 (APC-H7), CD56 (PerCP-Cy5.5), CD69 (BV711), TCRαβ (BUV563), TCRγδ (BUV395), CD8αβ (RY586), CD8αα (BB515), CD33 (BV480), CD49d/Itgα4 (PE-CF594), Itgβ7 (BUV737), CD103/ItgαE (BUV805), CD196/CCR6 (BUV661), and CD199/CCR9 (APC) (BD Biosciences, Franklin Lakes, NJ, USA); CD4 (BV421), HLA-DR (BV785), CD326/EpCAM (BV650), CD19 (PerCP-Cy5.5) and CD66b (PerCP-Cy5.5; BioLegend, San Diego, California, USA). The fixable Viability Stain 575V (BV570; BD Biosciences) was used to exclude dead cells from the analysis. Phenotypic analysis was performed by flow cytometry using a FACSymphony™ A5 Cell Analyzer, (BD Biosciences) and FlowJo software (Tree Star, Ashland, Oregon, USA).

### CellEngine Flow Cytometry Analysis

BD Diva FCS files were further exported and analyzed using the CellEngine software (CellCarta, Montreal, Canada). Briefly, the t-Distributed Stochastic Neighbour Embedding (t-SNE) and Self-Organizing Map (SOM) analysis were performed on CD3^+^ T-cells, CD3^+^CD4^+^ T-cells or CD3^+^CD8^+^ T-cells gated manually, as illustrated in Supplemental Figure 1. The following parameters were used for CD3^+^ T-cells: 25% subsampling, channels (TCRαβ, TCRγδ, CD4, CD8αα, CD8αβ, CD326, HLA-DR, CD69, CD196/CCR6, CD199/CCR9, CD49d/Itgα4, CD103/ItgαE, Itgβ7), t-SNE perplexity = 100, and number of nearest neighbors (k) = 400 (Figure 1). The following parameters were used for CD3^+^CD4^+^ T-cells and CD3^+^CD8^+^ T-cells: 50% subsampling, channels (TCRαβ, TCRγδ, CD326, HLA-DR, CD69, CD196/CCR6, CD199/CCR9, CD49d/Itgα4, CD103/ItgαE, Itgβ7), t-SNE perplexity = 100, and number of nearest neighbors (k) = 400 (Figure 2). The remaining parameters stayed as default settings and channels were rescaled equally (Figure 1-2).

**Figure 1:**
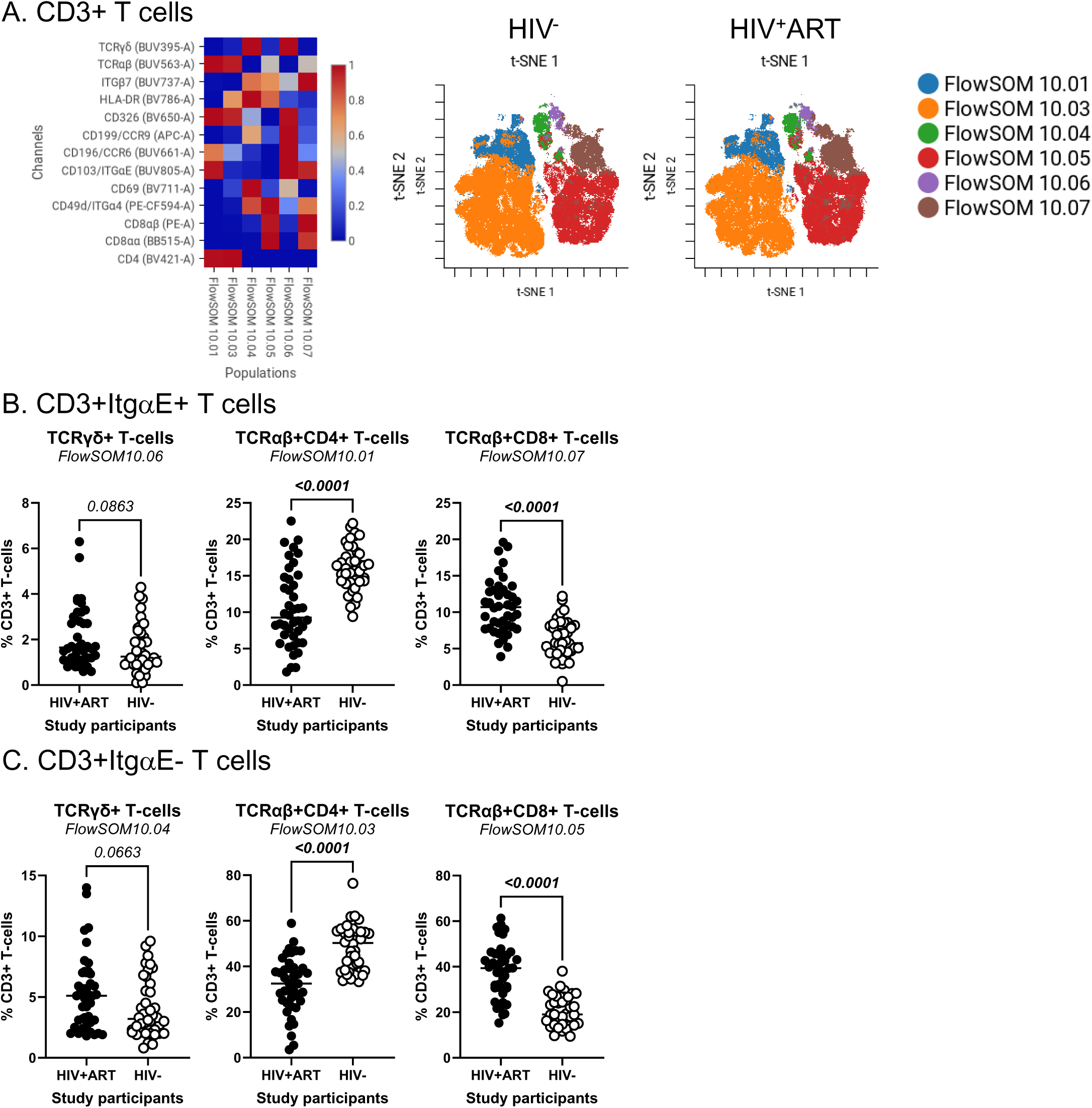
Identification of CD3^+^ T-cell subsets differentially expressed in ART-treated PWH *versus* HIV-uninfected participants. PBMCs from HIV+ART (n=42) and HIV-(n=40) participants were stained with a cocktail of fluorochrome-conjugated Abs for the identification of CD3^+^ T-cell subsets with differential expression of the T-cell lineage markers CD4, CD8αβ, CD8αα, TCRαβ, and TCRγδ, as well as the activation markers HLA-DR and CD69, the intestinal epithelial cell marker CD326, and the gut-homing chemokine receptors CD196/CCR6 and CD199/CCR9, and adhesion molecules CD49d/Itgα4, Itgβ7, CD103/ItgαE. The Fixable Viability Stain 575V was used to exclude dead cells from the analyses, as depicted in Supplemental Figure 1. The flow cytometry analysis was down sampled to 25% and analyzed using CellEngine with a perplexity = 100 and a number of nearest neighbours (k) = 400. Channels were rescaled equally on CellEngine. Shown are Heatmaps (left panel) and tSNE plots (middle and right panels) generated from supervised flow cytometry analysis of CD3^+^ T-cells **(A)**, as well as statistical analysis of the frequency of ItgαE^+^ **(B)** and ItgαE^-^ **(C)** CD3^+^ T-cell subsets with a TCRγδ^+^CD4^-^CD8^-^ (left panel), TCRαβ^+^CD4^+^ (middle panel) and TCRαβ^+^CD8^+^ T-cells (right panel). **(B-C)** Mann-Whitney p-values are indicated on the graphs.

**Figure 2:**
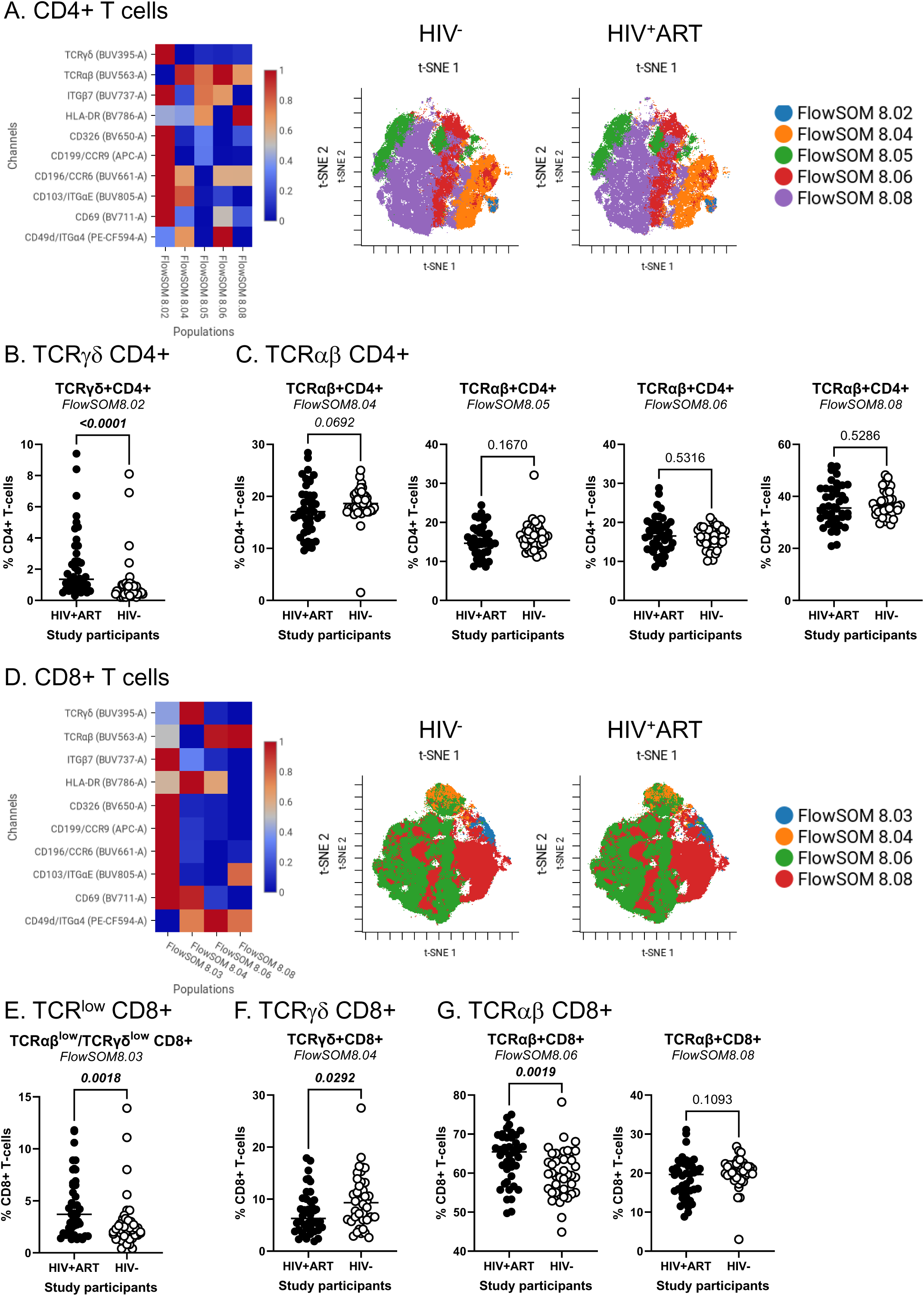
Identification of CD4^+^ and CD8^+^ T-cell subsets differentially expressed in ART-treated PWH *versus* HIV-uninfected participants. The flow cytometry staining of PBMCs from HIV+ART (n=42) and HIV-(n=40) participants was performed as detailed in Figure 1 legend. Analysis was down sampled to 50% and analyzed using CellEngine with a perplexity = 100 and a number of nearest neighbours (k) = 400. Channels were rescaled equally on CellEngine. **(A-G)** Shown are Heatmaps and tSNE plots generated from supervised flow cytometry data analysis of **(A-C)** CD3^+^CD4^+^ T-cells and (**D-G)** CD3^+^CD8^+^ T cells with differential expression of TCRγδ or TCRαβ, together with CD326, CCR6, CCR9, Itgα4, Itgβ7, and/or ItgαE. Shown are statistical analysis of the frequency of CD4^+^ **(B-C)** and CD8^+^ T-cell subsets **(E-G)** with TCRγδ^+^ **(B and F)** TCRγδ^low^TCRαβ^low^ **(E),** and TCRαβ^+^ phenotypes **(C and G)**. Mann-Whitney p-values are indicated on the graphs.

### Statistical analysis

The Shapiro-Wilk test was employed to evaluate whether continuous variables followed a normal distribution. Normally distributed continuous variables were described using mean and standard deviation (SD), whereas not normally distributed variables were described using median and interquartile range (IQR). Mann-Whitney test was used for comparing variables not normally distributed using the GraphPad Prism 10.3.0 software (Dotmatics, Boston, Massachusetts, USA). Fisher’s Exact Test was used to test if an association existed between two or three categorical variables. To identify novel immunological predictors of the HIV and CVD/TPV status and to determine the effect of potential confounding factors on the association between the circulating cell subsets and the HIV status and the presence or absence of coronary plaque within HIV^+^ individuals, we used a generalized linear model (GLM) regression as implemented in the R [54] package glmmTMB. For the prediction HIV status, the modeling was performed using a logistic regression (Supplemental Table 1). For the prediction CVD status and TPV values, the modeling was performed in two parts [55]. First, a logistic regression model was used to determine the association between covariates and the presence or absence of coronary plaques (The zero-inflation model) (Supplemental Table 2). Second, the association between covariates and the extent of plaque volumes among participants with the presence of coronary atherosclerosis was investigated by GLMs regression (The conditional model) (Supplemental Table 3). The confounding effect was investigated by the change in the linear model’s estimate for each subpopulation before and after the adjustment. We compared the crude (univariate) and the multiple regression models to study the contribution of the immune subset alone and in the presence of a sets of potential confounding factors (Supplemental Tables 1-3). A change in the estimated association measure of less than 10% identified small confounding bias, while a change superior to 30% identified large confounding bias [56, 57]. The p-value adjustment for the multiple testing hypothesis was performed according to the method of Benjamin and Hochberg [58], which controls the false discovery rate with adjusted p-value cutoffs of 0.05. Nominal p-values were adjusted within each T-cell subpopulation with a specific immune phenotype. Confounding modeling analysis with glmmTMB were performed using the statistical package R version 4.4.1.

## RESULTS

### Clinical and laboratory characteristics of the study participants

The HIV^+^ (n=42) and HIV^-^ (n=40) participants from the CHACS included in the cardiovascular imaging subgroup were categorized in TPV^+^ and TPV^-^, based on the presence or the absence of coronary atherosclerotic plaque. TPV was measured using coronary artery CCTA, as we previously reported [49, 51–53]. Detailed demographic and clinical information on study participants are included in Table 1. Briefly, HIV^+^ *versus* HIV^-^ participants were similar regarding age (median 54.1 *versus* 54.5 years old), male sex (95% *versus* 85%), and Framingham risk score (FRS; median 9 *versus* 10). HIV^+^ participants presented lower body mass index (BMI; median 24.1 *versus* 26.35 Kg/m^2^; p=0.0052). Smoking habits tended to be more prevalent among HIV^+^ compared to HIV^-^ participants, but differences did not reach statistical significance (p=0.1826). For TPV values, differences did not reach statistical significance between HIV^+^ and HIV^-^ when all participants where included (p=0.617). However, among TPV^+^ participants, TPV values were significantly higher in HIV^+^ compared to HIV^-^ groups (median 851 *versus* 182 mm^3^; p=0.0002). The HIV^+^ and HIV^-^ groups also presented with statistically similar levels of D-dimers (p=0.8127), fibrinogen (p=0.5735), HDL (p=0.1973), and triglycerides (p=0.2154); however, HIV^+^ participants had lower levels of LDL than the HIV^-^ group (median 2.55 *versus* 3.1 mmol/L; p=0.0452). The latter observation may be explained by the fact that a higher percentage of HIV^+^ participants were undergoing statin treatment (26% [11 of 42] compared to 12.5% [5 of 40]; p=0.0580).

The HIV^+^ participants grouped based on the presence (TPV^+^) or the absence (TPV^-^) of the atherosclerotic plaque were further analyzed for multiple clinical and laboratory parameters included in Table 2. Differences between the TPV^+^ and TPV^-^ groups did not reach statistical significance for age (p=0.6573), male sex (95.8% *versus* 93.7%; p>0.9999), BMI; (p=0.1320), FRS (p=0.1404), statin treatment (p=0.7275), plasma levels of D-dimer (p=0.3029), LDL (p=0.2972), HDL (p=0.7596), and triglycerides (p=0.2033), as well as the CMV co-infection status (p=0.5227), duration of HIV infection (p=0.2363), nadir CD4 counts (p=0.7889), and the frequency of CD4^+^ T-cells (p=0.2250). In contrast, the smoking habits tended to be higher (p=0.0666) among TPV^+^ in comparison to TPV^-^ ART-treated PWH participants, together with plasma fibrinogen levels (p=0.0827) and the duration of ART (p=0.0706).

### Immune profile alterations indicative of HIV and subclinical CVD status

Here, we sought to determine whether changes in the frequency and activation status of specific T-cell subsets with a gut-homing and/or gut-residency phenotype in the peripheral blood can be linked to the HIV status and the CVD risk in ART-treated PWH. To this aim, PBMCs from HIV^+^ and HIV^-^ CHACS participants, with (TPV^+^) and without (TPV^-^) subclinical atherosclerosis, were analyzed for the composition of the pool of hematopoietic CD45^+^CD3^+^ T-cells (Supplemental Figure 1). Among CD45^+^ cells, T-cells lacking the lineage markers CD19 (B cells), CD56 (NK cells), CD66b (neutrophils), and CD33 (myeloid cells), but expressing CD3 (T-cells) were further analyzed for the expression of CD4 (CD4^+^ T-cells) and CD8αβ (CD8^+^ T-cells) (Supplemental Figure 1). As expected, our results showed statistically significant decreased and increased numbers of CD4^+^ T-cells and CD8^+^ T-cells, respectively, in HIV^+^ compared to HIV^-^ participants, as well as decreased CD4/CD8 ratios (Supplemental Figure 2A). Aiming to investigate if there was a relationship between these parameters and the presence of plaque, we found that CD4^+^ and CD8^+^ T-cell numbers, as well as CD4/CD8 ratios, were similar regardless of the presence of plaque (Supplemental Figure 2B). For the HIV^-^ group, the CD4/CD8 ratios correlated positively with age and TPV values (Supplemental Figure 2C-D, right panels). These correlations were not observed in the HIV^+^ group (Supplemental Figure 2C-D, left panels), indicative of an increased CVD risk occurring regardless of aging in the context of HIV infection.

### CellEngine identification of CD4^+^ and CD8^+^ T-cell subsets with gut-homing phenotype and altered frequency and activation status during ART-treated HIV infection

Our group previously reported alterations in CD4^+^ T-cell phenotype and functions to be associated with the CVD status in ART-treated PWH [50, 59, 60]. To identify novel immunological predictor of HIV status and CVD risk, CellEngine analysis of polychromatic flow cytometry data was performed, upon staining with a set of T-cell lineage (CD45, CD3, CD4, CD8αα, CD8αβ, TCRαβ, TCRγδ), activation (HLA-DR), epithelial cell (EpCAM/CD326), and gut-homing/residency markers (CD69, CD196/CCR6, CD199/CCR9, CD49d/Itgα4, CD103/ItgαE, Itgβ7). The analysis of CD3^+^ T-cells revealed the presence of six major subsets, with differential expression of the T-cell lineage markers TCRαβ, TCRγδ, CD4, CD8αα, and/or CD8αβ, as well as the intraepithelial residency marker ItgαE [61], as follows: FlowSOM10.01 (TCRαβ^+^CD4^+^ItgαE^+^), FlowSOM10.03 (TCRαβ^+^CD4^+^ItgαE^-^), FlowSOM10.04 (TCRγδ^+^CD4^-^ CD8^-^ItgαE^-^), FlowSOM10.06 (TCRγδ^+^CD4^-^CD8^-^ItgαE^+^), FlowSOM10.05 (TCRαβ^+^CD8^+^ItgαE^-^), and FlowSOM10.07 (TCRαβ^+^CD8^+^ItgαE^+^) (Figure 1A, left heatmap). Of note, there was preferential expression of Itgβ7 on TCRγδ^+^ (FlowSOM10.04 and 10.06) and TCRαβ^+^CD8^+^ T-cells (FlowSOM10.05 and 10.07), CD326 on TCRαβ^+^CD4^+^ T-cells (FlowSOM10.01 and 10.03), CCR6 on CD4^+^ (FlowSOM10.01) and CD8^+^ (FlowSOM10.06) T-cell subsets, while CD69 and CCR9 were expressed at the highest levels on TCRγδ^+^ (FlowSOM10.04 and 10.06) T-cells (Figure 1A, left heatmap).

The frequency of these six clustered varied between HIV^+^ and HIV^-^ groups (Figure 1A, right t-SNE SOMs; Figure 1B-C). There was only a tendency for increased frequency of TCRγδ^+^ T-cell subsets expressing or not ItgαE (FlowSOM10.04 and 10.06) in HIV+ *versus* HIV-groups (Figure 1B-C, left panels). In contrast, differences reached statistical significance for decreased TCRαβ^+^CD4^+^ (FlowSOM10.01 and 10.03) (Figure 1B-C, middle panels) and increased TCRαβ^+^CD8^+^ (FlowSOM10.05 and 10.07) T-cell subsets (Figure 1B-C, right panels) in HIV^+^ *versus* HIV^-^ groups.

The CellEngine analysis was further performed on CD4^+^ and CD8^+^ T-cells, manually gated as illustrated in Supplemental Figure 1. The results depicted in Figure 2 reveal subtle T-cell subsets that varied in frequency and homing/activation phenotype between HIV^+^ and HIV^-^ groups for TCRγδ^+^ and TCRαβ^+^CD4^+^ (Figure 2A-C) and TCRαβ^+^CD8^+^ T-cells (Figure 2D-F). The highest expression of the gut/homing/residency markers Itgβ7, CD326, CD69, CCR6, CCR9 and ItgαE was observed on subsets of TCRγδ^+^CD4^+^ (FlowSOM8.02; Figure 2A) and CD8^+^ T-cells with relatively low TCRαβ and TCRγδ expression (FlowSOM8.03; Figure 2D), with their frequency being both increased in HIV^+^ *versus* HIV^-^ participants (Figure 2B and 2E). There was only a tendency for reduced frequency of TCRαβ^+^CD4^+^ T-cells (FlowSOM8.04; Figure 2C, left panel). Other significant differences were observed for TCRγδ^+^CD8^+^ (FlowSOM8.03) and TCRαβ^+^CD8^+^ T-cells (FlowSOM8.06), with their frequency being decreased and increased, respectively, in HIV^+^ *versus* HIV^-^ participants (Figure 2F and 2G, left panel).

In addition to the expected depletion of TCRαβ^+^CD4^+^ T-cells and expansion of TCRαβ^+^CD8^+^ T-cells in the peripheral blood of PWH, these analyses reveal an altered composition of these circulating T-cell pools during ART-treated HIV-infection, with the abundance of subsets with a gut-homing/residency profile in HIV^+^ *versus* HIV^-^ participants. Such an accumulation may reflect the aberrant mobilization of key sentinel cells from the gut into the periphery, likely as a consequence of gut barrier impairment.

### Increased frequencies of CD4^+^ T-cells with a gut-homing/residency CD326^+^CD69^+^CCR6^+^ phenotype in HIV^+^ *versus* HIV^-^ participants

The CellEngine flow cytometry analysis (Figure 1A and 2A) brought our attention to the presence of a fraction of TCRαβ^+^ CD4^+^ T-cells expressing the epithelial cell marker EpCAM/CD326 [62–64] and the gut-homing marker CCR6 (enriched in FlowSOM8.08, Figure 2A). This presence was further confirmed by manual gating in Figure 3A, with CD326^+^CD4^+^ T-cells being more abundant in HIV^+^ compared to HIV^-^ participants (p=0.001) (Figure 3B). Compared to total CD4^+^ T-cells, the CD326^+^CD4^+^ T-cell subset of HIV^+^ participants distinguished from total CD4^+^ T-cells by a statistically significant increased expression of CD69 (p=0.008), CCR6 (p<0.0001) and ItgαE (p<0.0001), a tendency for increased CCR9 expression (p=0.0686), and a decreased expression of Itgβ7 (p=0.0015) and Itgα4 (p<0.0001), with no significant differences for HLA-DR (Figure 3C). The same phenotypic differences between total and CD326^+^CD4^+^ T-cells were also observed among HIV^-^ participants (data not shown). Of particular interest, CD326^+^CD4^+^ T-cells of HIV^+^ *versus* HIV^-^ participants expressed at higher frequency CD69 (p=0.0002) and CCR6 (p=0.0389) (Figure 3D), with now significant differences for the other markers (data not shown). These results point to the enriched presence in the circulation of HIV^+^ *versus* HIV^-^ participants of an atypical subset of TCRαβ^+^CD4^+^ T-cells expressing the epithelial marker CD326 and overexpressing the gut homing/residency markers CD69 and CCR6.

**Figure 3:**
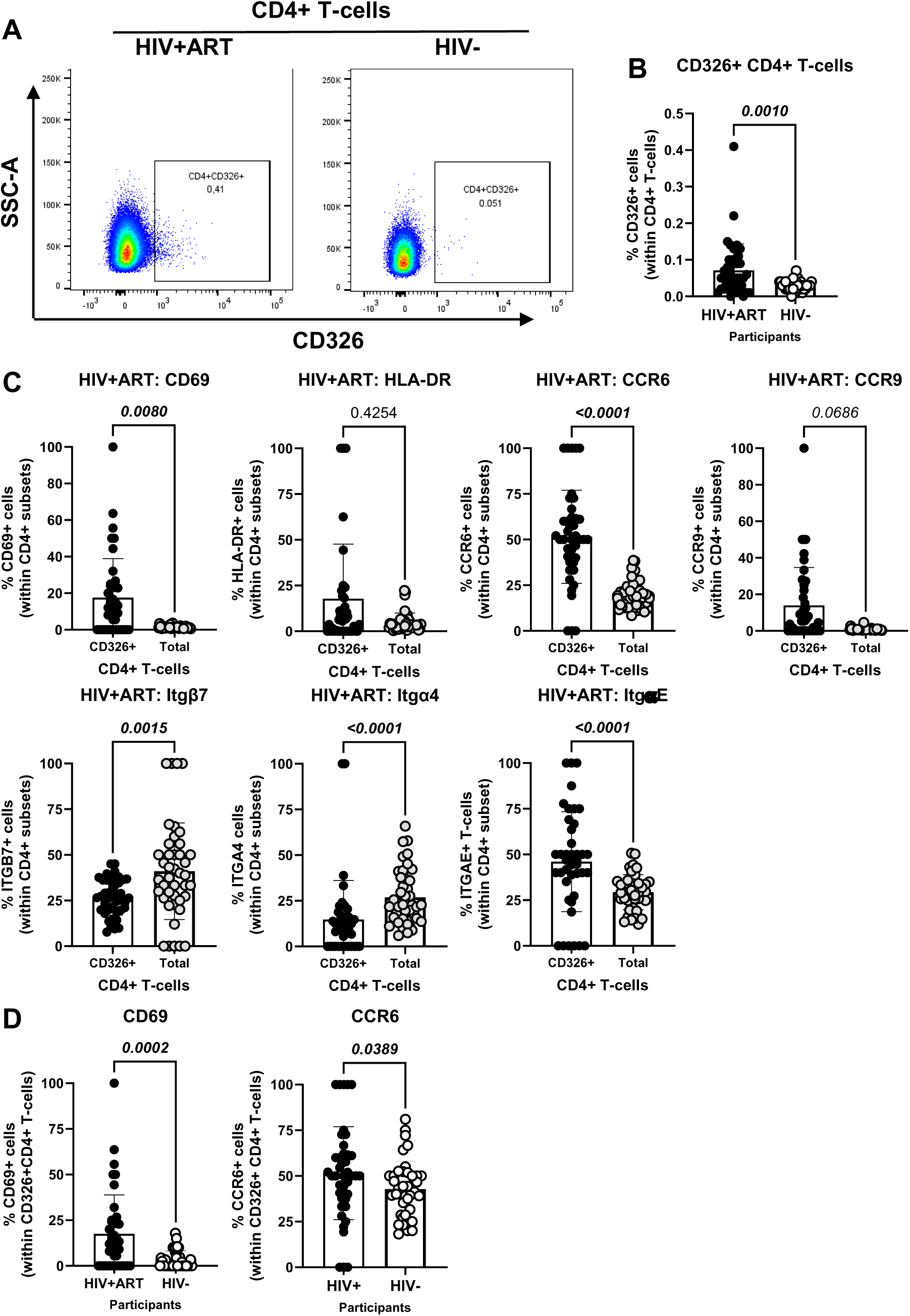
Frequency and phenotype of peripheral blood CD326^+^ CD4^+^ T-cells in HIV^+^ *versus* HIV^-^ participants. The flow cytometry staining of PBMCs from HIV+ART (n=42) and HIV^-^ (n=40) participants was performed as detailed in Figure 1 legend. The flow cytometry analysis was performed by manual gating using the FlowJo software, as depicted in Supplemental Figure 1. Shown are the representative gating strategy for the identification of CD326^+^ CD4^+^ T-cells **(A)** and statistical analysis of their frequency **(B)** in HIV+ART *versus* HIV^-^ participants; and **(C)** the expression of activation stained with a cocktail of molecules (CD69 and HLA-DR), and gut-homing chemokine receptors (CCR6 and CCR9) and integrins (Itgβ7, Itgβ4 and ItgαE) on CD326^+^ CD4^+^ T-cells of HIV^+^ participants, as well as differences in CD69 and CCR6 expression between CD326^+^ CD4^+^ T-cells of HIV^+^ *versus* HIV^-^ participants **(D)**. Mann-Whitney p-values are indicated on the graphs.

### Increased frequencies of CCR6^+^ItgB7^-^ CD4^+^ T-cells in the peripheral blood of HIV^+^ *versus* **HIV^-^ participants**

The CellEngine flow cytometry analysis pointed to the presence of four TCRαβ^+^CD4+ T-cells subsets with differential expression of the gut-homing markers CCR6 and Itgβ7 (Figure 2A). We further performed manual gating on FlowJo to investigate the frequency and phenotype (i.e., CD69, HLA-DR, ItgαE, CCR9) of CD4^+^ T-cells expressing or not CCR6 and/or Itgβ7 (Figure 4). We found no significant differences in the frequency of CCR6^+^Itgβ7^+^ CD4^+^ T-cells (enriched in FlowSOM8.06, Figure 2A) in HIV^+^ compared to HIV^-^ participants (Figure 4A-B) and no significant differences in the expression of CD69, HLA-DR, and ItgαE on CCR6^+^Itgβ7^+^ CD4^+^ T-cells between HIV^+^ and HIV^-^ participants, with CCR9 expression being slightly lower in HIV^+^ participants (Figure 4D). Contrariwise, the frequency of CCR6^+^Itgβ7^-^ CD4^+^ T-cells (enriched in FlowSOM8.08, Figure 2A) was higher in HIV^+^ compared to HIV^-^ participants (Figure 4A and C), with the expression of CD69 and HLA-DR being equivalent in both groups, and the ItgαE and CCR9 expression levels being slightly lower in HIV^+^ *versus* HIV-participants (Figure 4D). Of note, the expression of ItgαE and CCR9 was markedly lower in CCR6^+^ItgB7^-^ compared to CCR6^+^ItgB7^+^ CD4^+^ T-cells, indicative of an altered gut-homing potential of these cells in ART-treated PWH.

**Figure 4:**
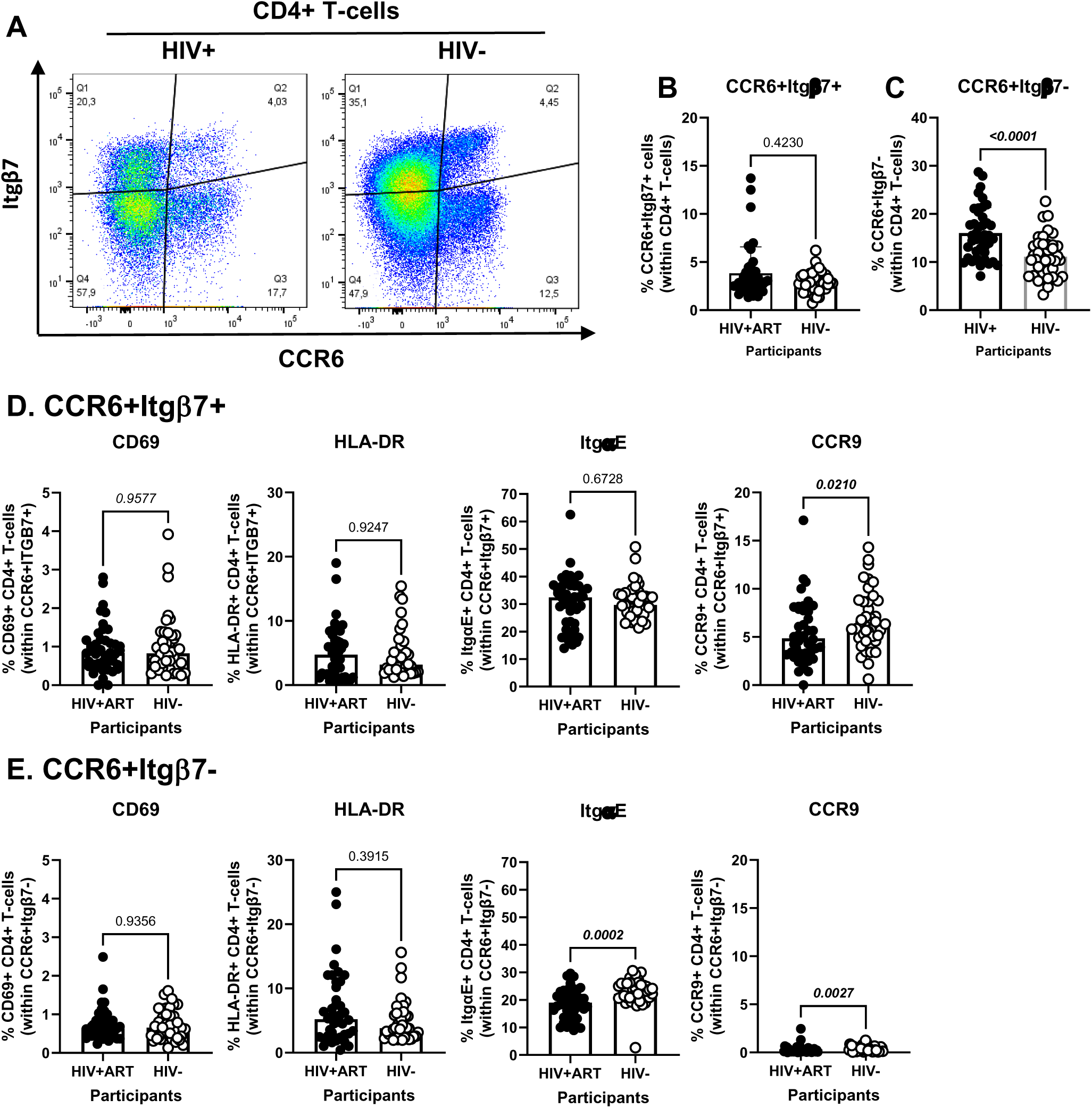
Frequency and phenotype of peripheral blood CD4^+^ T-cells expressing CCR6 and/or Itgβ7 in ART-treated PWH and HIV-uninfected participants. **(A-E)** PBMCs from HIV^+^ART (n=42) and HIV^-^ (n=40) participants were stained with a cocktail of fluorochrome-conjugated Abs, as detailed in Figure 1 legend. Cell frequencies and phenotypes were analyzed by flow cytometry manual gating (BD Fortessa and FlowJo), with CD4^+^ T-cells identified as in Supplemental Table 1. Shown is the gating strategy for the identification of CD4^+^ T-cells expressing CCR6 and/or Itgβ7 **(A)**, the frequencies of CD4^+^ T-cells with a CCR6^+^Itgβ7^+^ **(B)** and CCR6^+^Itgβ7^-^ **(C)** phenotypes in HIV+ART *versus* HIV^-^ participants; and the characterization of the expression of CD69, HLA-DR, ItgαE, and CCR9, on circulating CCR6^+^Itgβ7^+^ **(D)** and CCR6^+^Itgβ7^-^ **(E)** CD4^+^ T-cells in HIV+ART *versus* HIV^-^ participants. **(B-E)** Mann-Whitney p-values are indicated on the graphs.

### Decreased frequencies and gut-homing potential of CD8^+^ IEL-like T-cells in the peripheral blood of HIV^+^ *versus* HIV^-^ participants

Another element of our hypothesis of observing gut immune alterations in peripheral blood was related to the presence of circulating IELs. Using well-established markers in the literature [40, 42–46], we used manual gating on FlowJo to identify subsets of circulating CD4^+^ (CD3^+^CD4^+^Itgβ7^+^ItgαE^+^TCRαβ^+^TCRγδ^-^CD8αβ^-^) and CD8^+^ (CD3^+^CD4^-^Itgβ7^+^ItgαE^+^TCRαβ^+^TCRγδ^-^CD8αα^+^CD8αβ^-^) with IEL-like phenotypes (Figures 5-6). We observed that CD4^+^ IEL-like T-cells (Figure 5A) did not differ in frequency between HIV+ and HIV-groups (Figure 5B), but distinguished from their total CD4^+^ T-cell counterparts by superior expression of CD69 (p<0.0001), CCR6 (p<0.0001) and CCR9 (p<0.0001), with CCR6 being expressed on the majority of CD4^+^ IEL-like T-cells (Figure 5C), indicative of their imprinting for gut-homing. Of note, CD4^+^ IEL-like T-cells from HIV^+^ *versus* HIV^-^ groups expressed similar levels of CD69 and CCR6 but lower CCR9 expression (Figure 5D), indicative of a deficit in their CCR9-mediated homing. Further, our results show a decreased frequency of CD8^+^ IEL-like T-cells (Figure 6A) in HIV^+^ compared to HIV^-^ participants (p=0.0014) (Figure 6B), together with similar levels of CD69 and CCR6, but decreased CCR9 expression (p=0.0067) in cells from HIV^+^ *versus* HIV^-^ groups (Figure 6C). Together these results reveal the presence of circulating TCRαβ^+^CD4^+^ and TCRαβ^+^CD8^+^ T-cell subsets with IEL-like features, with CD8^+^ IEL-like cells being decreased in frequency and exhibiting a deficient CCR9-mediated gut-homing potential in association with the HIV status.

**Figure 5:**
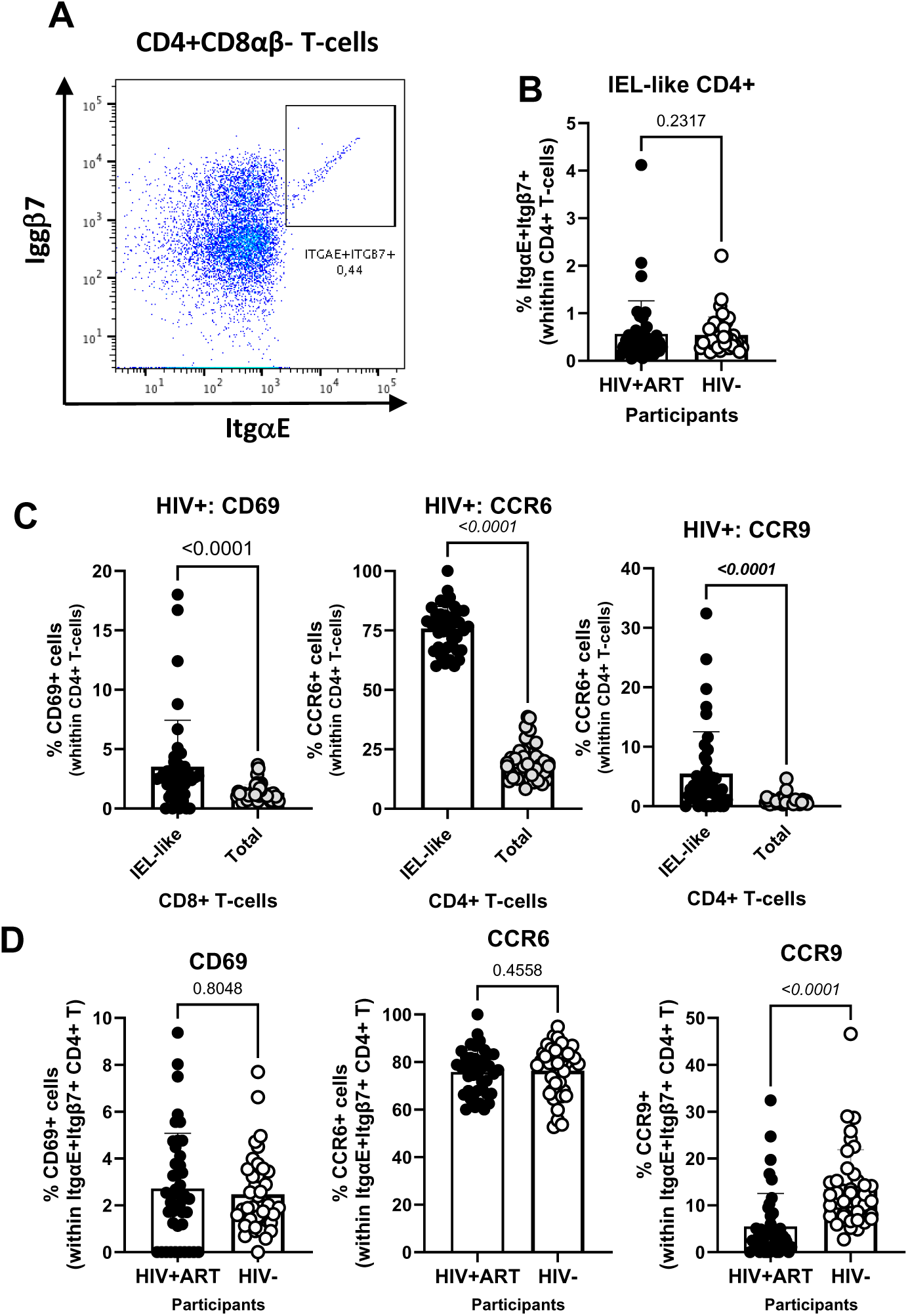
Frequency and phenotype of peripheral blood IEL-like CD4^+^ T-cells in ART-treated PWH *versus* HIV-participants. PBMCs from HIV+ART (n=42) and HIV-(n=40) participants were stained with a cocktail of fluorochrome-conjugated Abs, as in Figure 1 legend. The Fixable Viability Dye 575V was used to exclude dead cells from the analyses. Cell frequencies and phenotypes were analyzed by flow cytometry (BD Fortessa and FlowJo). The flow cytometry analysis was performed by manual gating, as depicted in Supplemental Figure 1. Shown is the gating strategy for the identification of CD4^+^ T-cells (CD45^+^CD3^+^CD4^+^TCRαβ^+^) with an IEL-like phenotype (Itgβ7^+^CD103/ItgαE^+^) **(A)**, the analysis of their frequency in HIV^+^ *versus* HIV^-^ participants **(B)**; as well as the expression of CD69, CCR6 and CCR9 on IEL-like *versus* total CD4^+^ T-cells **(C)**, and differences in the phenotype of IEL-like CD4^+^ T-cells between HIV+ART and HIV^-^ participants **(D)**. Mann-Whitney p-values are indicated on the graphs **(B-D)**.

**Figure 6:**
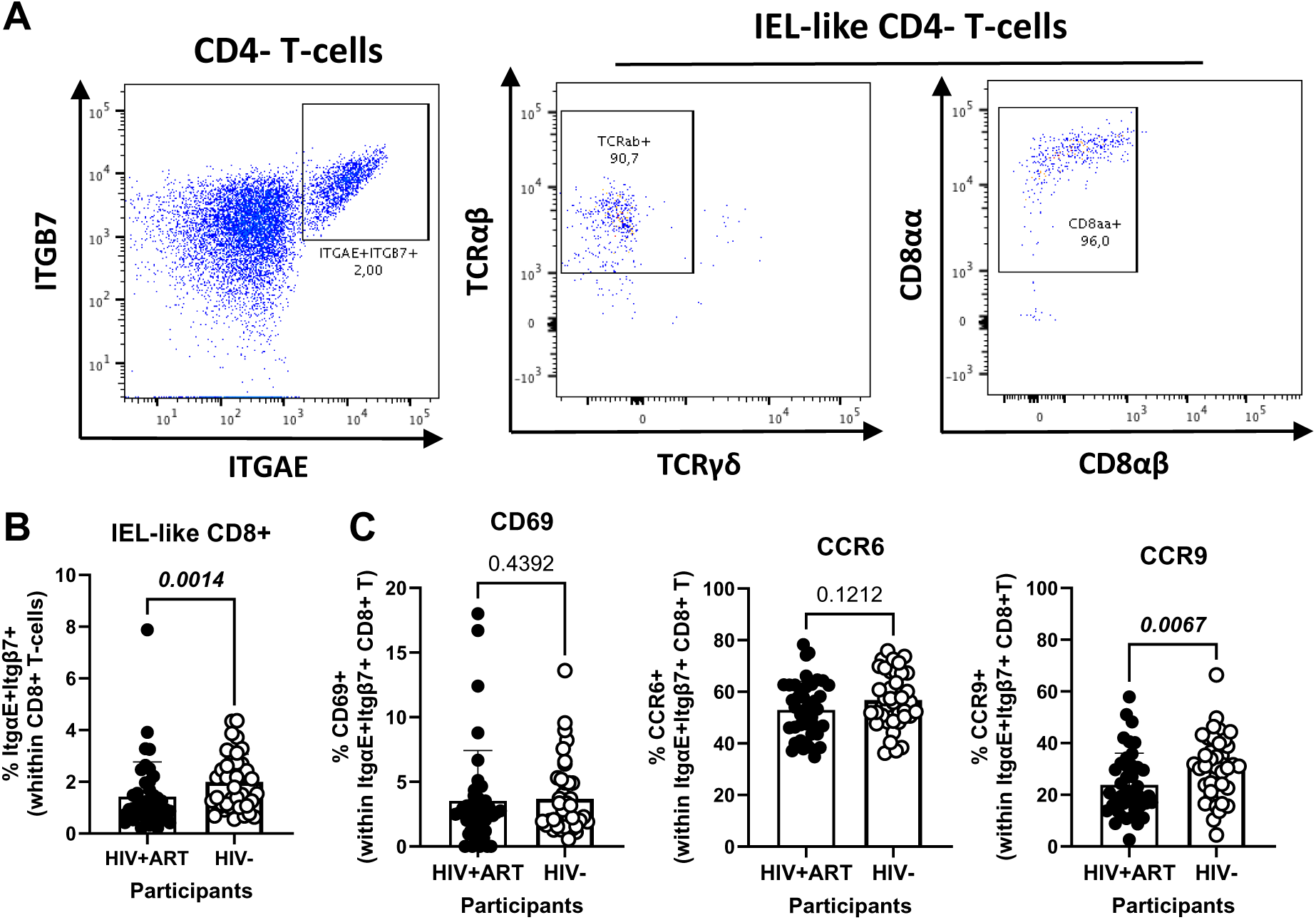
Frequency and phenotype of peripheral blood CD8+ IEL-like T-cells in HIV+ *versus* HIV-participants. **(A-F)** PBMCs from HIV+ART (n=42) and HIV-(n=40) participants were stained with a cocktail of fluorochrome-conjugated Abs, as detailed in Figure 1 legend, to allow the identification of CD8^+^ T-cells (CD45^+^CD3^+^CD4^-^TCRαβ^+^) with an IEL-like phenotype (Itgβ7^+^CD103/ItgαE^+^CD8αα^+^) and the analysis of their CD69, CD196/CCR6, and CD199/CCR9 expression. Shown are representative dot plots for the identification of Itgβ7^+^CD103/ItgαE^+^ CD4-T-cells (left panel) and their co-expression of TCRαβ/TCRαα (middle panel) and CD8αα/CD8αβ (right panel) **(A)** and the statistical analysis of the frequency of circulating IEL-like CD8^+^ T-cells in HIV^+^ *versus* HIV^-^ participants **(B)**; as well as the expression of CD69, CCR6 and CCR9 on IEL-like CD8^+^ T-cells from HIV^+^ *versus* HIV^-^ participants **(C)**. Mann-Whitney p-values are indicated on the graphs **(B-C)**.

### Multivariate logistic regression analyses identify immune subpopulations that predict the HIV status and the CVD risk in ART-treated PWH

To identify novel immunological predictors the HIV status and CVD risk, we performed a multivariate logistic regression analysis considering all studied immunological subpopulations and the clinical information listed in Supplemental File 1, with statistical results detailed in Supplemental Tables 1-3. To identify predictors of the HIV status, the basic logistic regression model, which only included the immune subsets (Crude Model), was evaluated against multivariate logistic regression models that adjusted for different potential confounding factors linked to cardiovascular health (*i.e.,* BMI, CVD status, LDL) and HIV status (*i.e.,* CD4 counts, CD8 counts, CD4/CD8 ratios, time on ART, time since HIV diagnosis), separately or all confounding factors together (ALL1 and ALL 2), as detailed in Supplemental Table 1. Across all models, the frequency of the following TCRαβ^+^CD3^+^ T-cell subpopulations was positively associated with the HIV status CD326^+^CD4^+^, CD326^+^CD69^+^CD4^+^, CCR6^+^Itgβ7^-^CD4^+^, CD8^+^FlowSOM8.03 and CD8^+^FlowSOM10.07 T-cells, while the CD4^+^CCR9^+^ IEL-like, CD4^+^ItgαE^+^FlowSOM10.01 and CD4^+^ItgαE-FlowSOM10.03 were identified as negative predictors (Supplemental Table 1, Supplemental Figure 3). Other subpopulations significant in the crude model but that lost statistical significance with the HIV status when adjustments were made for HIV-specific cofounding factors, included TCRγδ^+^FlowSOM10.05 and TCRαβ^+^CD8^+^FlowSOM08.06 (Supplemental Figure 3).

A similar multivariate logistic regression analysis failed to identify T-cell subpopulations associated with CVD in the HIV^+^ group, in the crude model or upon univariate adjustment for D-dimer, smoking status and FRS, as well as for HIV-specific confounding factors (*i.e.,* CD4 counts, CD8 counts, CD4/CD8 ratios, time on ART, time since HIV diagnosis) (Supplemental Table 2). The only exception was for CD326^+^CD69^+^ CD4^+^ T cells that were identified as weak positive predictors of the CVD status, with marginal significance reached in the crude model only (p=0.04711) (Supplemental Table 2). Nevertheless, when a similar strategy was used to identify immunological predictors of the TPV values in the ART-treated PWH group (Supplemental Table 3), multiple TCRαβ^+^CD3^+^ T-cell subpopulations were identified as positive and negative predictors (Figure 7A). Particularly, CD4^+^ItgαE^+^FlowSOM10.01 and CD4^+^ItgαE^-^FlowSOM10.03 where negatively correlated with the TPV values (Figure 7B), while CD8^+^ItgαE^+^FlowSOM10.07 and CD8^+^ItgαE^-^FlowSOM10.05 (Figure 7C), together with CCR6^+^CD4^+^ and CCR6^+^Itgβ7^-^CD4^+^, and at a lower extend CCR6^+^Itgβ7^+^CD4^+^ T-cells (Figure 7D), where identified as positive predictors in the crude model and upon adjustment to all or the majority of CVD and HIV-specific confounding factors (Figure 7A, Supplemental Table 3).

**Figure 7:**
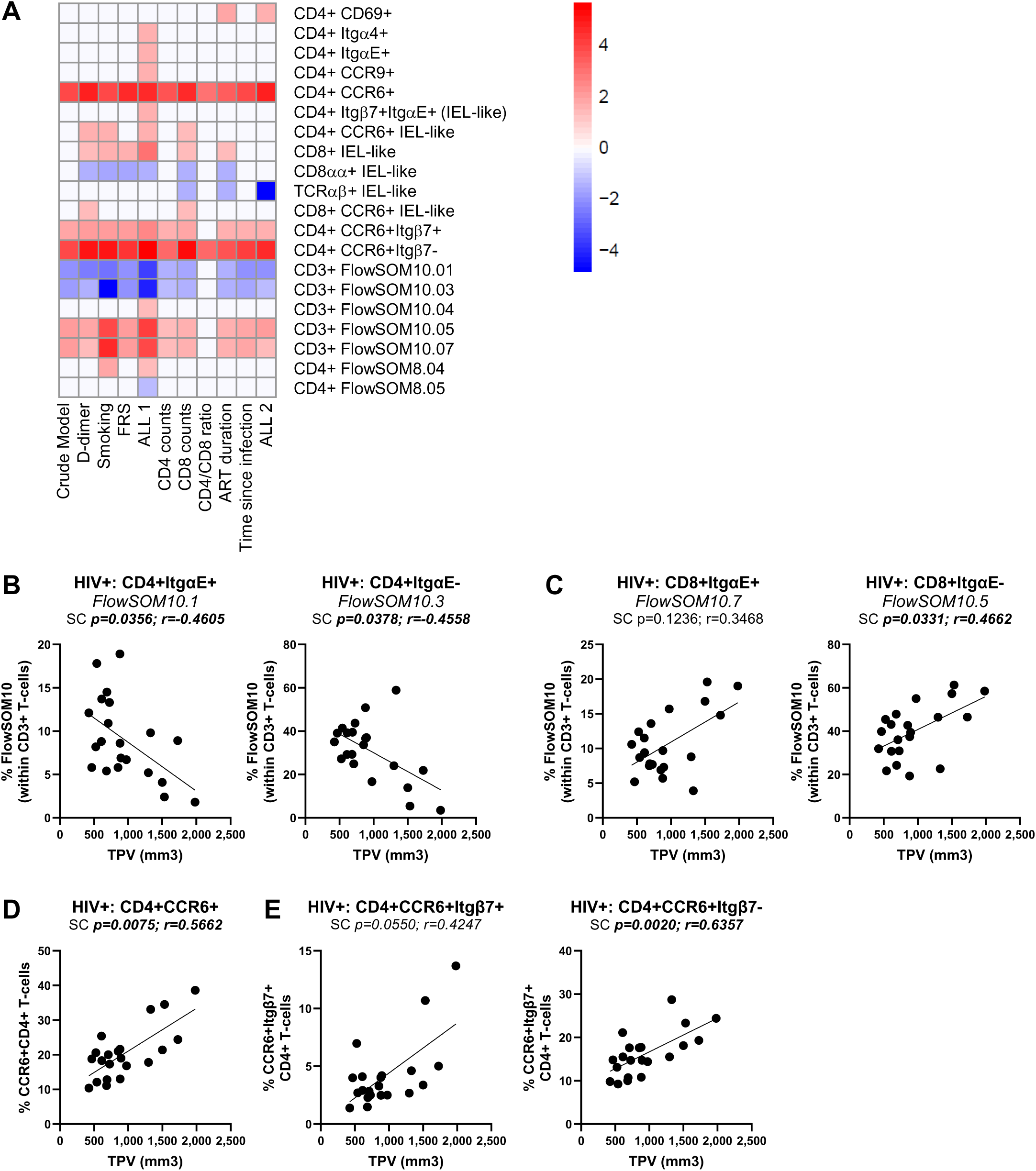
Multivariate regression model identified novel immunological predictors of subclinical atherosclerosis in ART-treated PWH. The heatmap illustrates the subsets of T-cells that were identified in Supplemental Table 3 as significant predictors of TPV values in the crude model or upon adjusting for D-dimer, smoking status, FRS, ALL 1 (D-dimer, smoking status, FRS), as well as CD4 counts, CD8 counts, CD4/CD8 ratios, time on ART, time since HIV diagnosis, and ALL 2 (CD4 counts, CD8 counts, CD4/CD8 ratios, time on ART, time since HIV diagnosis). Adjusted p-values (-log10) are indicated as a red and blue gradient for positively (Sign=1) and negatively associated predictors (Sign=-1), respectively. **(A).** Shown are individual illustrations of the correlations TPV values and the frequency of CD4^+^ **(B)** and CD8^+^ T-cells **(C)** with an ItgαE^+^ (left panels) or ItgαE^-^ phenotypes (right panels), as well as CD4^+^ T-cells with a CCR6^+^ phenotype **(D)** expressing or not Itgβ7 **(E)**. Spearman correlation p and r values are indicated on the graphs **(B-D)**.

Together, these results point to major changes in the frequency, activation status, and gut-homing/residency potential of specific CD4^+^ and CD8^+^ T-cell subsets circulating in the peripheral blood that robustly predict the HIV status and the presence of the subclinical atherosclerotic plaque. Such peculiar T-cell subsets may represent new peripheral blood markers for evaluating and therapeutically managing the CVD risk in ART-treated PWH.

## Discussion

In this study, we identified alterations in the composition of the pool of circulating CD3^+^ T-cells expressing gut-homing/residency markers as novel predictors of HIV-associated coronary atherosclerosis among ART-treated CHACS participants. This study was inspired by the concept of liquid biopsy used in oncology [65–67] and the evidence that inflammatory conditions in the intestines are associated with the unusual circulation in the blood of intestinal epithelial cells and other tissue-resident immune cells [68–70]. Our major findings point to ***i)*** aberrant circulation of CD326^+^CD4^+^ T-cells, TCRγδ^+^, and TCRαβ^+^ cells with IEL-like phenotypes in the peripheral blood of HIV^+^ *versus* HIV^-^ participants and ***ii)*** a positive association between TPV and the abundance of CCR6^+^CD4^+^ T-cells expressing or not the gut-homing marker Itgβ7, subsets, previously identified by our group as being highly permissive to HIV infection [71]. These results are consistent with intestinal barrier alterations persisting during HIV-1 infection [15, 16, 28–30] and a documented link between HIV reservoir size in CD4^+^ T-cells and subclinical CVD [72].

PBMCs from n=82 participants of the CHACS were analyzed, including 42 ART-treated PWH and 40 individuals without HIV. Both groups were further categorized based on the presence/absence of coronary atherosclerotic plaque. HIV^+^ individuals in the TPV^+^ group showed an increased burden of coronary atherosclerosis compared to HIV^-^TPV^+^ participants, consistent with previous research by our group [50, 53]. The analysis of the HIV^+^ group revealed that TPV^+^ compared to TPV^-^ participants had similar FRS and plasma D-dimer levels, two documented predictors of cardiovascular risk [73, 74]. The TPV^+^ and TPV^-^ HIV^+^ participants also showed a similar time since HIV infection, duration of ART, nadir CD4 counts, and current CD4 percentages. Interestingly, age and TPV correlated with the CD4/CD8 ratio in HIV^-^ but not in HIV^+^ individuals, showing opposite trends between the two groups. These clinical similarities between TPV^+^ and TPV^-^ HIV^+^ participants prompted us to search for new immunological predictors of the subclinical atherosclerosis in ART-treated PWH. Thus, we performed polychromatic flow cytometry analysis using a large panel of T-cell lineage (CD45, CD3, CD4, CD8αα, CD8αβ, TCRαβ, TCRγδ), epithelial cell (EpCAM/CD326), activation (HLA-DR), and gut-homing/residency markers (CD69, CD196/CCR6, CD199/CCR9, CD49d/Itgα4, CD103/ItgαE, Itgβ7). This allowed an in depth characterization of circulating T-cell subsets differentially expressed in CHACS participants with known HIV and TPV status.

A noteworthy finding of our study was the increased abundance of a subset of TCRαβ^+^CD4^+^ T-cells expressing the epithelial cell adhesion molecule (EpCAM)/CD326 in HIV^+^ compared to HIV^-^ participants. CD326 plays a crucial role in establishing a functional intestinal barrier [63, 64, 75], is expressed on sigmoid colon epithelial cells [76], as well as circulating tumor epithelial cells used as markers in the context of liquid biopsies [64, 65, 77]. CD4^+^ T-cells might acquire CD326 expression upon interaction with epithelial cells through trogocytosis, a process that involves the transfer of surface molecules and membrane fragments from one cell to another, thereby granting recipient cells new functionalities or the ability to eliminate donor cells [78, 79]. Such mechanism has been reported in the context of HIV-1 infection [80] and may reflect the history of cell-to-cell interactions *in vivo*. Nevertheless, the intrinsic expression of CD326 mRNA in T-cells require further evaluation at transcriptional level.

Compared to total CD4^+^ T-cells, CD326^+^CD4^+^ T-cells expressed a CD69^+^ItgαE^+^CCR6^+^CCR9^+^ phenotype. CD69 is an early T-cell activation marker and a promotor of T-cell retention in tissues [13], with its expression being associated with T-cell activation in HIV-1 infection [81–83]. By comparing the phenotype of the CD326^+^CD4^+^ T-cells to total CD4^+^ T-cells within HIV^+^ participants, we observed increased expression of CD103/ItgαE, a hallmark of gut residency, considering the interaction between CD103/ItgαE on T-cells and E-cadherin on intestinal epithelial cells [40, 84]. Moreover, we have also observed increased CCR6 expression in CD326^+^CD4^+^ T-cells. Previous research by our group and others indicated that CCR6-expressing T-cells are more susceptible to HIV-1 and harbor higher amounts of integrated HIV-DNA in ART-treated PHW [28–30]. Furthermore, CD326^+^CD4^+^ compared to total CD4^+^ T-cells from HIV^+^ individuals were characterized by heightened expression of CCR9, with the CCR9/CCL25 axis being key for gut-homing, a process altered during HIV infection [16, 37, 85]. Of note, CCR9 is associated with inflammatory conditions and its knockdown can reduce CVD progression [86]. This is in line with the aberrant accumulation of CCR9^+^ T-cells in the blood of ART-treated PWH, as a consequence of their impaired recruitment into the gut due to a deficit of mucosal CCL25 expression [37]. All together, these results support the idea that the presence of CD326^+^CD4^+^ T-cells in the circulation may reflect alterations in the intestinal barrier integrity during HIV disease progression.

Both CD4^+^ and CD8^+^ T-cells expressing CD103/ItgαE and CD69 are affected during HIV-1 infection [16, 48, 84, 85]. Of note, CellEngine analysis also depicted high CD326 expression on TCRγδ^+^CD4^-^CD8^-^ItgαE^+^ (FlowSOM10.06) and TCRαβ^low^TCRγδ^low^CD4^-^CD8^-^ItgαE^+^ (FlowSOM8.03) T-cells, subsets that also showed increased abundance in HIV^+^ *versus* HIV^-^participants. Of note, unconventional T-cells that co-expressed TCRαβ and TCRγδ have been described before [87, 88]. Like Th17 cells, TCRγδ^+^T-cells are crucial for maintaining the gut barrier integrity [45] and are significantly perturbed during HIV-1 infection, even during effective ART [89–91]. The aberrant presence of different CD326^+^ T-cell subsets in the circulation may reflect the state intestinal barrier impairment, well documented in the context of ART-treated HIV infection and associated with the CVD risk.

Another interesting finding of our study was the increased frequency of CCR6^+^Itgβ7^-^ CD4^+^ T-cells in HIV^+^ *versus* HIV^-^ individuals, without significant changes on the frequency of CCR6^+^Itgβ7^+^CD4^+^T-cells. Of note, CCR6^+^Itgβ7^-^ and CCR6^+^Itgβ7^+^ CD4^+^ T-cells from HIV^+^ *versus* HIV^-^ CHACS participants expressed lower levels of CCR9 and ItgαE, respectively, pointing to their altered gut-homing potential. While memory CD4^+^ T-cells expressing both integrin α4β7 and CCR6 are inclined towards a Th17 lineage and are more vulnerable to HIV, it is primarily the expression of CCR6, rather than integrin α4β7 that reflects this susceptibility [71, 92, 93]. The relative contribution of CCR6^+^Itgβ7^-^ *versus* CCR6^+^Itgβ7^+^ CD4^+^ T-cells to HIV reservoir persistence during ART requires further investigations. Nevetherless, our group and others have previously reported that CCR6 expression is a marker of HIV reservoirs [93–95]. Other research documented the presence of integrated HIV-DNA in CD4^+^CD103^+^CD69^+^CCR6^low^ T-cells infiltrating the gut of PWH [47, 48]. Of particular notice, we also identified a fraction of CD4^+^ T-cells with IEL-like (ItgαE^+^Itgβ7^+^) and gut-homing/residence phenotype (CD69^+^CCR6^+^CCR9^+^). Although their frequency did not differ inside the pool of CD4^+^ T-cells of HIV^+^ *versus* HIV^-^, their role as HIV reservoirs during ART deserves further investigations.

Our study revealed a decreased frequency of CD8^+^ T-cells with an IEL-like phenotype (ItgαE^+^Itgβ7^+^TCRαβ^+^TCRγδ^-^CD8αα^+^CD8αβ^-^) in ART-treated PWH compared to HIV^-^ counterparts, together with a reduced CCR9 expression, again indicative of their impaired intraepithelial homing capacity. The cell phenotype was characterized by TCRαβ and CD8αα expression, which reveal that they are “natural” IELs, distinct from “induced” IELs that primarily express CD4 or CD8αβ and derive from conventional TCRαβ^+^T-cells of peripheral lymphoid tissues. Natural IELs originate from precursor cells that have undergone development in the thymus and can migrate to the intestinal epithelium without the need for a cognate antigen [40, 42, 46]. Initially discovered in mice, the presence of TCRαβ^+^CD8αα^+^ IELs in humans is still debatable [42], although a low frequency (<1%) is documented in humans [46]. While their TCRαβ^+^CD8αβ^+^ and TCRγδ^+^CD8αα^+^ counterparts perform distinct functions within the immune system, TCRαβ^+^CD8αα^+^ IELs are particularly noted for their potential suppressive/regulatory contributions, as well as similarities antiviral NK cells [42, 46]. Due to the lack of description of these IELs in human studies, research into their role in HIV infection is limited. Studies reported a significant reduction in the population of both CD4^+^ and CD4^+^CD8αα^+^ IELs within the intestinal epithelium during the SIV/HIV infection [96]. Other research mentions alterations in subsets that simultaneously expressed CD8αα and αβ [97]. Notably, in this work, the subset we examined did not exhibit co-expression of these markers. Therefore, to our knowledge, this is the first study documenting the diminution of TCRαβ^+^CD8αα^+^ IELs in the peripheral blood of PWH. Since IELs are key in targeting virus-infected IECs, via perforin, granzyme B, and IFN-γ and can perform CTL functions and secrete a broad spectrum of interferons, including types I, II, and III, supporting the antiviral state of the epithelium [43, 98], we speculate that their reduced frequency is associated with impaired antiviral responses during HIV infection.

Finally, multivariate logistic regression analyses linked the changes in the composition of circulating T-cell subsets expressing gut-homing/residency markers to the HIV and CVD status, as well as coronary atherosclerosis plaque values. Of particular relevance, we identified ItgαE^+^CD8^+^, ItgαE^-^CD8^+^, CCR6^+^CD4^+^, and CCR6^+^Itgβ7^-^CD4^+^ T-cell subsets as positive predictors, and the ItgαE^+^CD4^+^ and ItgαE^-^CD4^+^ T-cell subsets as negative predictors of TPV values in crude models or upon adjustment for HIV (*i.e.,* CD4 counts, CD8 counts, CD4/CD8 rations, time on ART, time since infection) and CVD (*i.e.,* D-dimers, smoking status, FRS) confounding factors.

Our study has several limitations. The study was performed in majority on male participants and the validity of these results on female participants remains to be determined. The predictive value of the identified cellular markers need to be validated in large scale immune monitoring studies during specific interventions designed to decrease the CVD risk in ART-treated PWH (*e.g.,* statins) [99]. The study is descriptive and future mechanistic studies, as well as longitudinal investigations, are needed to link events occurring in the gut to the alterations observed in the pool of circulating T-cells and demonstrated the direct contribution the identified subsets to the occurrence of CVD in ART-treated PWH.

In conclusion, our results reveal alterations in the composition of the pool of circulating T-cell subsets with a gut-homing/residency and activated phenotype that are predictive of the HIV status and the coronary atherosclerotic plaque presence and volume. The increased frequency of CD326^+^CD4^+^ T-cells and decreased numbers of natural CD8^+^ IELs in the peripheral blood suggest ongoing gut barrier disruption, which may predispose ART-treated PWH to increased CVD risk, as well documented [8, 15, 24, 25]. This study also identifies a specific immune cell phenotype, CCR6^+^Itgβ7^-^CD4^+^ T-cells, positively associated with increased atherosclerotic plaque burden in PWH. Our results provide a compelling rationale for further investigation into targeted interventions that address not only viral suppression but also the unique immune mucosal dysregulation in ART-treated PWH, reflected in the peripheral blood, for reducing the CVD onset and mortality and improving long-term health outcomes in an aging PWH community.

## AUTHORS’ CONTRIBUTIONS

EMG performed the experiments, analyzed the results, prepared the figures, and wrote the manuscript. JD helped with PBMC processing for flow cytometry analysis. REC helped with color compensation in Symphony. TWS generated preliminary data on circulating CD326^+^ IECs. MN provided expertise with IELs. CC-L acquired and interpreted the cardiovascular imaging data. AF-M performed the multivariate statistical analysis. MEF, MD and CT provided protocols, access to human samples, and expertise with CVD. J-PR provided access to human samples. PA designed the study, analyzed the results, designed the figures, and revised the manuscript. All authors revised and approved the manuscript.

## ACKNOWLEDGEMENTS

The authors thank Philippe St-Onge and Gael Dulude (Flow Cytometry Core Facility, CHUM-Research Centre) for expert technical support with polychromatic flow cytometry sorting; Dr Anita L. Ray (CellCarta, Montréal, Qc, Canada) for training in CellEngine Analysis; Olfa Debbeche for managing the NLC3 Core Facility of the CHUM-Research Centre; Dr. Annie Chamberland, Stéphanie Matte, Mohamed Sylla for blood processing and managing the cell biobank; Marc Messier-Peet for providing clinical information and managing the CHACS database; and Mario Legault for his help with ethical approvals and informed consents. The authors address a special thanks to all ART-treated PWH and HIV-uninfected study participants for their crucial contribution to this work.

## SUPPLEMENTAL FIGURE LEGENDS

**Supplemental Figure 1 (related to Figures 1-6): General gating strategy for flow cytometry analysis of PBMCs of study participants.** PBMCs from HIV^+^ART (n=22) and HIV^-^ (n=20) participants with known CVD status, measured as the total plaque volume (TPV; mm^3^) of the coronary artery atherosclerosis, were stained with the Fixable Viability Stain 575V for dead cell exclusion and with a cocktail of abs that allowed the exclusion of other lineage cells (CD33, CD56, CD19 and CD66b), and the identification of CD45^+^ hematopoietic cells. Among CD45^+^ cells, CD3^+^ T-cells were further differentiated into CD4^+^ and CD8αβ^+^ T-cells.

**Supplemental Figure 2 (related to Figure 1): Frequency of CD4+ and CD8+ T-cells in HIV+ *versus* HIV-participants relative to age and subclinical CVD.** PBMCs from study participants were identified as per Supplemental Figure 1. Shown are **(A)** the frequency of CD4^+^ T-cells, CD8^+^ T-cells and the CD4/CD8 ratio relative to the HIV status; **(B)** the characterization of CD4^+^ T-cells, CD8^+^ T-cells and CD4/CD8 ratio relative to the presence of subclinical CVD; **(C)** the correlation between age and the CD4/CD8 ratio in HIV^+^ **(left panel)** and HIV^-^ **(right panel)** participants; and **(D)** the correlation between TPV and the CD4/CD8 ratio in HIV^+^ **(left panel)** and HIV^-^ **(right panel)** participants. Mann-Whitney p-values, as well as Spearman correlation p and r values, are indicated on the graphs.

**Supplemental Figure 3 (related to Supplemental Table 1): Multivariate regression model identified novel immunological predictors of HIV-1 status.** Heatmap illustrate the subsets of T-cells that were identified in Supplemental Table 1 as statistically significant predictors of the HIV-1 status in the crude model, or upon adjusting for parameters identified as statistically different between HIV+ART and HIV-groups (Table 1), as follows: BMI, CVD status, LDL, ALL 1 (BMI, CVD status, LDL), as well as CD4 counts, CD8 counts, CD4/CD8 ratios, and ALL 2 (CD4 counts, CD8 counts, CD4/CD8 ratios). Adjusted p-values (-log10) are indicated as a red and blue gradient for positively (Sign=1) and negatively associated predictors (Sign=-1), respectively.

