## Supplemental Figures 1-3 for "Alterations in Circulating T-Cell Subsets with Gut-Homing/Residency Phenotypes Predict HIV-1 Status and Subclinical Atherosclerosis"

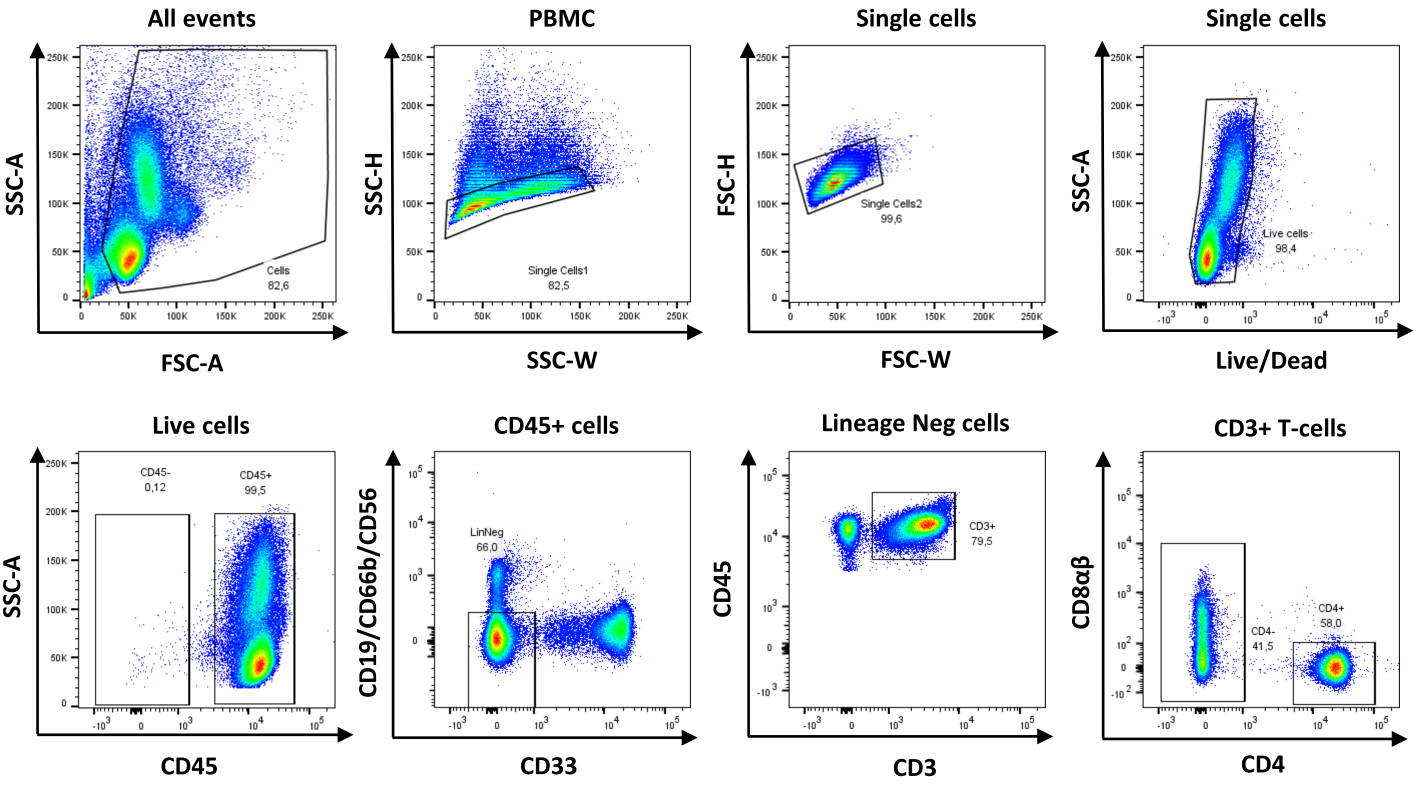


Moreira Gabriel/Dias et al., Supplemental Figure 1
Relative to Figures 1-6

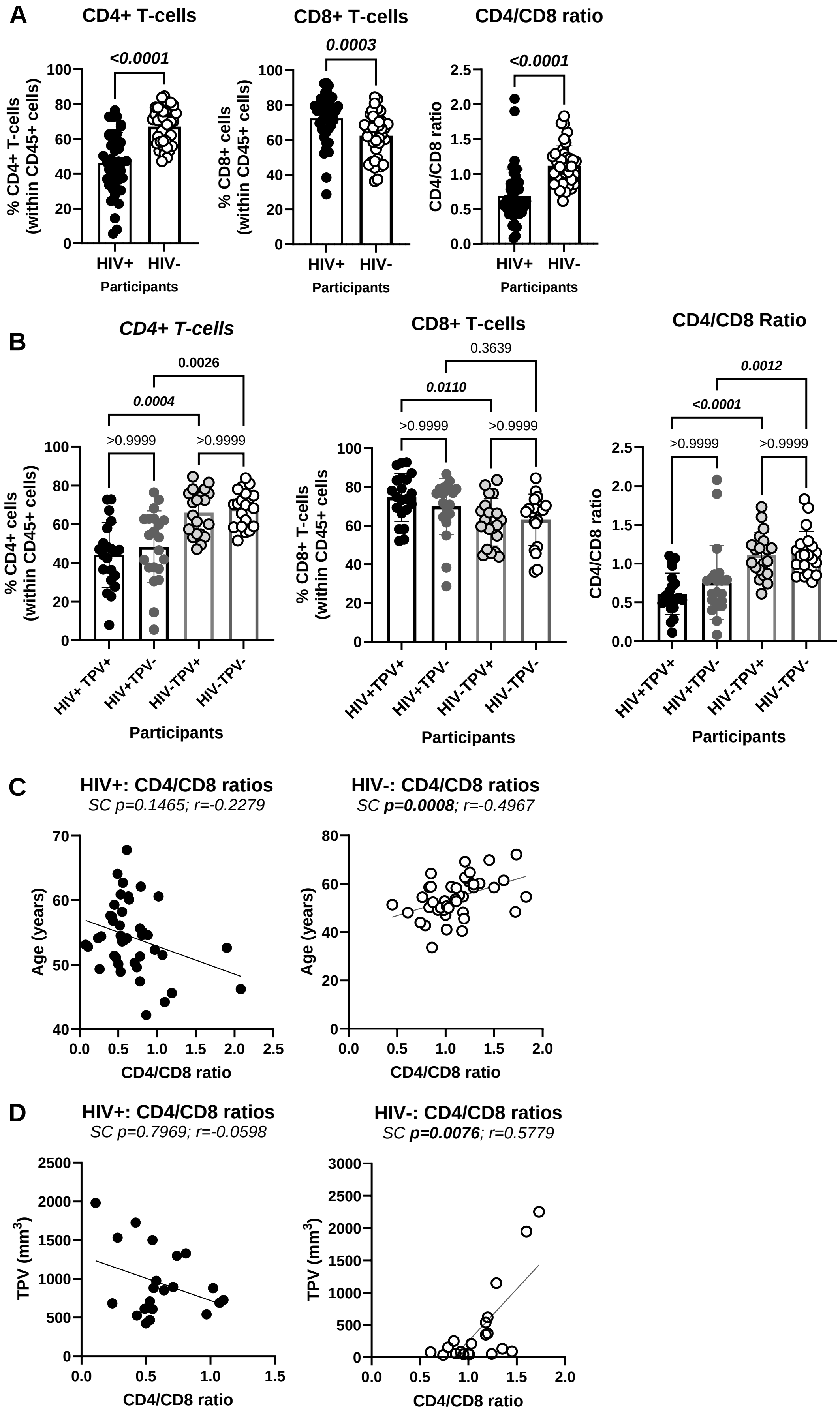


Moreira Gabriel/Dias et al., Supplemental Figure 2
Relative to Figure 1

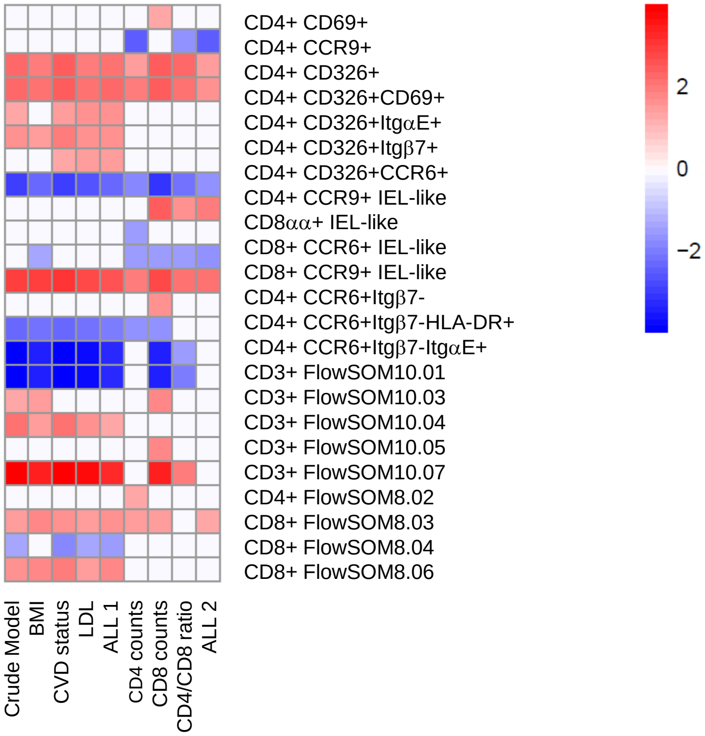


Moreira Gabriel/Dias et al., Supplemental Figure 3
Relative to Supplemental Table 1
